# Complex effects of kinase localization revealed by compartment-specific regulation of protein kinase A activity

**DOI:** 10.1101/2021.04.08.439038

**Authors:** Rebecca LaCroix, Benjamin Lin, Andre Levchenko

## Abstract

Kinase activity in signaling networks frequently depends on regulatory subunits that can both inhibit activity by interacting with the catalytic subunits and target the kinase to distinct molecular partners and subcellular compartments. Here, using a new synthetic molecular interaction system, we show that translocation of a regulatory subunit of the protein kinase A (PKA-R) to the plasma membrane has a paradoxical effect on the membrane kinase activity. It can both enhance it at lower translocation levels, even in the absence of signaling inputs, and inhibit it at higher translocation levels, suggesting its role as a linker that can both couple and decouple signaling processes in a concentration-dependent manner. We further demonstrate that superposition of gradients of PKA-R abundance across single cells can control the directionality of cell migration, reversing it at high enough input levels. Thus complex *in vivo* patterns of PKA-R localization can drive complex phenotypes, including cell migration.

## Introduction

In intracellular signal transduction, the information is encoded in molecular interactions involving recognition of diverse substrates by specific enzymes. These interactions are facilitated by large sets of adapter and scaffold proteins linking the activated enzymes to substrates within specific subcellular compartments, controlling dynamic modifications of protein localization and local concentrations of effector molecules (Langeberg & Scott, 2015). Furthermore, the enzymes, such as the diverse and abundant kinases involved in cell signaling and other functions, frequently contain covalently linked regulatory and catalytic subunits, with the regulatory subunits controlling both the activity of the enzyme and its interactions with other enzymes and substrates. However, the enzymes belonging to the family of protein kinase A (PKA) serine-threonine kinases do not follow this covalent linkage rule. Instead, a PKA molecule is a complex of two catalytic subunits (PKA-C) and a regulatory subunit dimer (PKA-R) that can dissociate following binding of two cyclic AMP (cAMP) molecules to each of the PKA-R subunits (Taylor, Zhang, Steichen, Keshwani, & Kornev, 2013). This intricate organization of the kinase complex is further complicated by dynamically shifting subcellular pools of cAMP, tethering of the kinase and its substrates to a large family of differentially localized A-kinase anchoring proteins (AKAPs), localized phosphatase and phosphodiesterase activity, and the intrinsic inhibitory function of PKA-R (Baillie, 2009; Wong & Scott, 2004; Zhang et al., 2012). What emerges is a picture of structural and functional complexity of PKA signaling that is still incompletely understood in spite of decades of research and analysis.

Given the complexity of intermolecular interactions, the stoichiometry of multi-molecular complexes can have profound effects on the outcome of signaling processes. A particularly striking effect is observed if a linker protein, such as a scaffold molecule can vary in its abundance. It has been shown, e.g., for the MAPK signaling pathways, that a scaffold protein can enhance the pathway activity at an optimal level but can also inhibit it if its concentration exceeds the optimum (Chapman & Asthagiri, 2009; Levchenko, Bruck, & Sternberg, 2000). This ‘combinatorial inhibition’ effect (Good, Zalatan, & Lim, 2011) suggests that variation of the relative abundance of the pathway components can modulate the pathway activity even if the input levels do not change. Given the structural complexity of PKA signaling, it is not clear whether and how the relative abundance of various signaling pathway components in different subcellular compartments may modulate the signaling outcomes. More specifically, it is not clear if PKA-R subunits would serve purely as inhibitors of PKA signaling (as expected due to their intrinsic inhibitory role, relieved only in the presence of high cAMP concentrations) or would potentially elevate the kinase activity by enhancing localization of the PKA holoenzyme to subcellular areas with increased signaling inputs.

Compartmentalized PKA signaling is important in a number of contexts including glucose homeostasis, cardiomyocyte contractility, cell cycle regulation, and cell migration (Howe, 2004; Langeberg & Scott, 2005; Mauban, O’Donnell, Warrier, Manni, & Bond, 2009; Wong & Scott, 2004). In regulating cell migration, PKA is known to positively affect the activity of some cytoskeletal regulators (e.g., Rac1, Cdc42, α4β1 integrin) and negatively affect others (e.g., RhoA). Additionally, both inhibition and activation of PKA have been shown to have inhibitory effects on cell migration (Howe, 2004). As a result, PKA’s role in regulation of cell migration is still unclear. Several studies utilizing FRET biosensors have identified gradients of PKA activity in migrating cells, with relatively high PKA activity at the cell front and relatively low activity at the rear, suggesting that spatial control of the kinase is involved in this process (Lim et al., 2008; Paulucci-Holthauzen et al., 2009; Tkachenko et al., 2011). Furthermore, the regulatory but not the catalytic subunit has been shown to be enriched in the pseudopods of cells in culture, suggesting that regulatory subunit localization may play a role in the regulation of cell migration (Howe, Baldor, & Hogan, 2005).

Since PKA-R mediates anchoring of PKA to diverse intracellular locations, in large part due to its interactions with AKAPs, it is particularly important to explore whether and how its abundance at specific subcellular locations modulates the output of signaling activity both under the basal conditions and in response to specific stimulation. This analysis can benefit from a tool that can permit acute localization of PKA-R to a predefined cell location in the absence of direct pathway stimulation. We developed such a tool based on chemically-inducible dimerization with the small, cell-permeable molecule rapamycin (Banaszynski, Liu, & Wandless, 2005). Rapamycin induces dimerization of two small, intracellularly transduced molecular components: FK506-binding protein (FKBP) and the FKPB-rapamycin domain (FRB), which can be tethered to proteins of interest as well as specific subcellular compartments. This technique has been used to study biochemical activity of different proteins (Chu et al., 2014; Dagliyan et al., 2017; Dagliyan et al., 2013; Inoue, Heo, Grimley, Wandless, & Meyer, 2005; Karginov, Ding, Kota, Dokholyan, & Hahn, 2010). Previously, we demonstrated that this dimerization strategy, when combined with the use of a microfluidic device controlling spatial rapamycin distribution, can be used to study biomolecular systems controlling cell polarity and migration (Lin et al., 2012). For the current study, we have tethered FKBP to fluorescently labeled PKA-R, while anchoring FRB to the plasma membrane (PM), to enable dynamically controlled localization of PKA-R to the PM in a rapamycin-dependent fashion. Using this tool, we find that PKA-R can have unexpectedly complex regulatory effects on the activity and function of PKA at the PM, elucidating the potential role of PM PKA-R localization in normal and pathological cell function.

## Results

### Design and characterization of an inducible PKA-R translocation system

To induce rapid and spatially-controlled translocation of PKA-R to the plasma membrane, we utilized a rapamycin-based dimerization strategy combined with in-dish or in-chip control of rapamycin concentration similar to the strategy we previously used to control Rac1 function (Lin et al., 2012). We linked one binding partner of rapamycin, FKBP, to a fluorescently tagged PKAR-IIβ (PKAR-FKBP-FP) and the other, FRB, to the membrane-targeting sequence of Lyn kinase (Lyn_11_-FRB, also referred to as Lyn-FRB throughout the text) (Figure 1A). Two color variants of the PKA-R construct were created to facilitate co-imaging with fluorescent biosensors and dyes (Figure 1B and Figure 1 – figure supplement 1). In addition to our transient expression vectors, lentiviral Gateway expression vectors for PKAR-FKBP-FP and Lyn-FRB were created and integrated into the genome of HeLa cells for ease of experimentation. We will refer to these cell lines as HeLa PFM (mCherry variant) and HeLa PFY (YFP variant) respectively.

**Figure 1.**
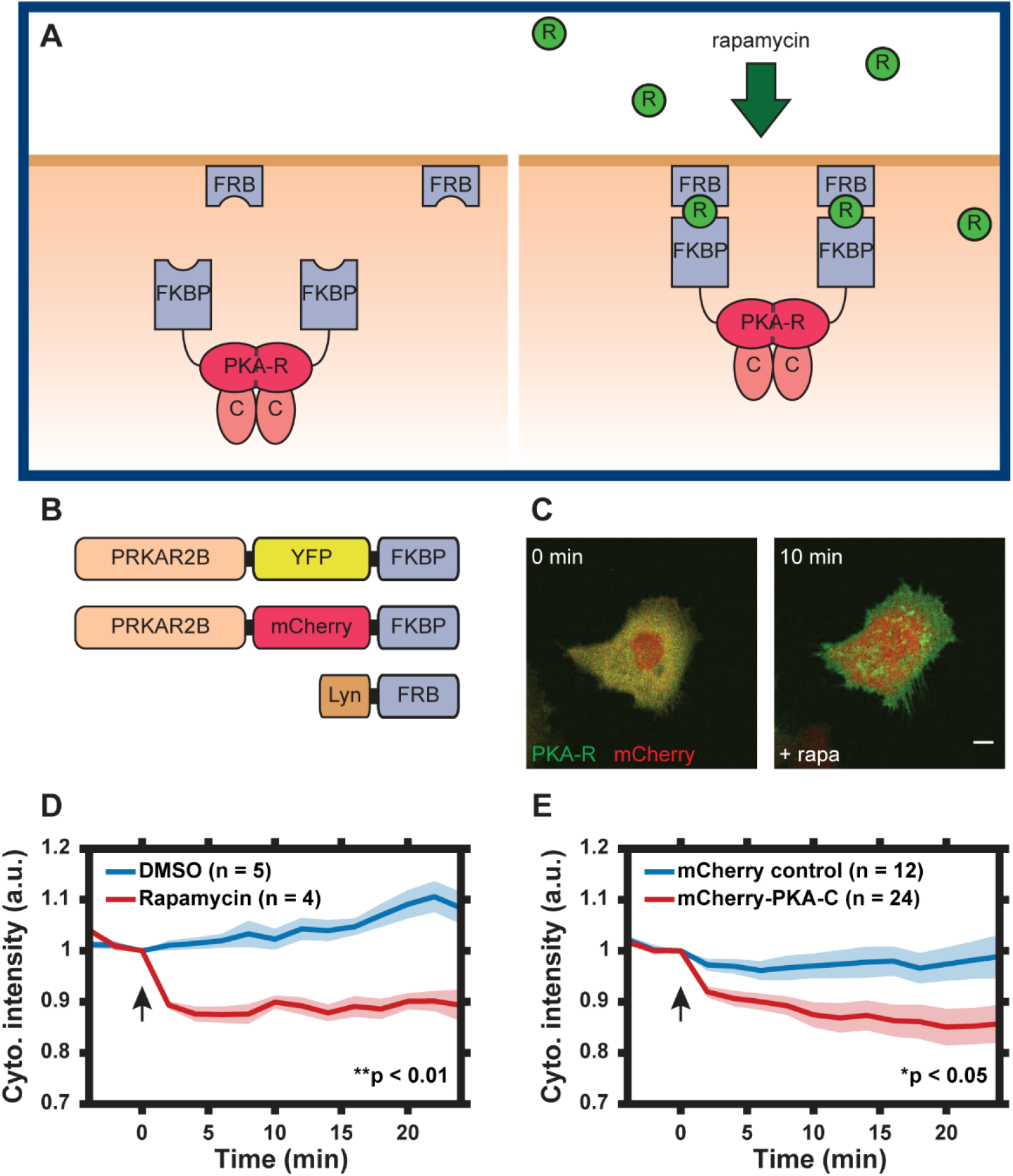
Design of PKA-R translocation system. (A) Schematic of PKA-R translocation system. Rapamycin induces heterodimerization of FKBP and FRB, resulting in translocation of PKA-R to the plasma membrane. [FKBP = FK506-binding protein, FRB = FKBP-rapamycin binding domain, R = rapamycin, PKA-R = PKA regulatory subunit, C = PKA catalytic subunit] (B) DNA construct design. Two versions of recombinant PKA-R were created with different fluorescent labels. (C) PKA-R localization (green) at 0 and 10 mins after addition of 100 nM rapamycin. Scale bar, 10 µm. mCherry protein (red) co-expressed for visualization. (D) PKA-R translocation quantified as cytoplasmic intensity drop in YFP channel following addition of DMSO or 100 nM rapamycin. p = 0.0039 at t = 24 min post-rapamycin addition; two-tailed Student’s *t*-test. (E) PKA-C translocation quantified as a cytoplasmic intensity drop in mCherry channel following addition of 100 nM rapamycin. p = 0.037 at t = 24 min post-rapamycin addition; two-tailed Student’s *t*-test. Graphs display the mean of each data set with SEM indicated by shaded region. Number of cells in each data set is as indicated in the figure. Data is the result of 1 (D) and 3 (E) independent experiments, respectively. Arrows indicate the timing of drug addition.

To test translocation of PKA-R to the membrane, HeLa cells transiently expressing the YFP-tagged translocation system were treated with 100 nM rapamycin and imaged for 30 minutes. Membrane translocation was measured as a decrease in the cytoplasmic intensity in the YFP channel. Addition of rapamycin resulted in a rapid PKA-R translocation to the PM evaluated as a significant decrease of the cytoplasmic fluorescence intensity over the first three minutes of stimulation (Figures 1C and 1D). No significant translocation occurred after this initial period. To determine whether PKA-C translocated to the membrane along with the exogenous PKA-R, we co-transfected HeLa PFY cells with the constructs encoding either recombinant mCherry protein or mCherry-tagged PKA-C. Following addition of rapamycin, PKA-C indeed translocated to the cell membrane whereas mCherry alone underwent no significant change in localization, demonstrating that our synthetic system can bring the intact PKA holoenzyme to the plasma membrane (Figure 1E and Videos 1 and 2).

### Characterization of cell response to PKA-R translocation

The dissociation of PKA-C from PKA-R is commonly seen as a relief of PKA-C inhibition within the holoenzyme, with the regulatory subunit thus treated as a negative regulator of the kinase activity. To test whether membrane translocation of PKA-R would indeed inhibit the basal PKA activity in this compartment, we transfected HeLa PFM cells with Lyn-AKAR4, a membrane-bound FRET probe for PKA activity (Depry, Allen, & Zhang, 2011). The dynamic range of the intracellular Lyn-AKAR4 responses in this cell line was determined to be 20.9% ± 2.0% (n = 12) [mean ± standard error of the mean (SEM) (n = number of cells)] following cell treatment with a combination of an adenylyl cyclase activator forskolin (Fsk, 50 μM) and a competitive non-selective phosphodiesterase inhibitor 3-isobutyl-1-methylxanthine (IBMX, 100 μM) to maximally increase the intracellular cAMP levels (Figure 2 – figure supplement 1). Surprisingly, we found that rapamycin-induced PKA-R translation alone (without any additional stimulation) was able to induce a significant increase in PKA activity, which was transient in some cells and sustained in others (Figures 2A and 2B). To determine whether the variability in the concentration of PKA-R could account for this cell-cell variation of response, we examined the dependence of the maximum levels of PKA activity on the estimated PKA-R abundance, using mCherry fluorescence intensity as a proxy. Interestingly, we observed that the increase in activity was greatest for cells with intermediate PKA-R concentrations, decreasing when PKA-R was either higher or lower than this optimal level (Figure 2C). Furthermore, in the cells with the highest PKA-R expression, PKA activity, following the initial rise, later not only decreased vs. the maximum, but dropped below the basal level, indicating active inhibition of the PKA activity (Figures 2D and 2E). Notably, even in the cells in which the PKA activity was inhibited versus the basal levels, this activity transiently increased, suggesting that the PKA-R translocation was gradual, and thus first reached activating levels at the PM, but ultimately exceeded these levels and reached inhibitory concentrations.

**Figure 2.**
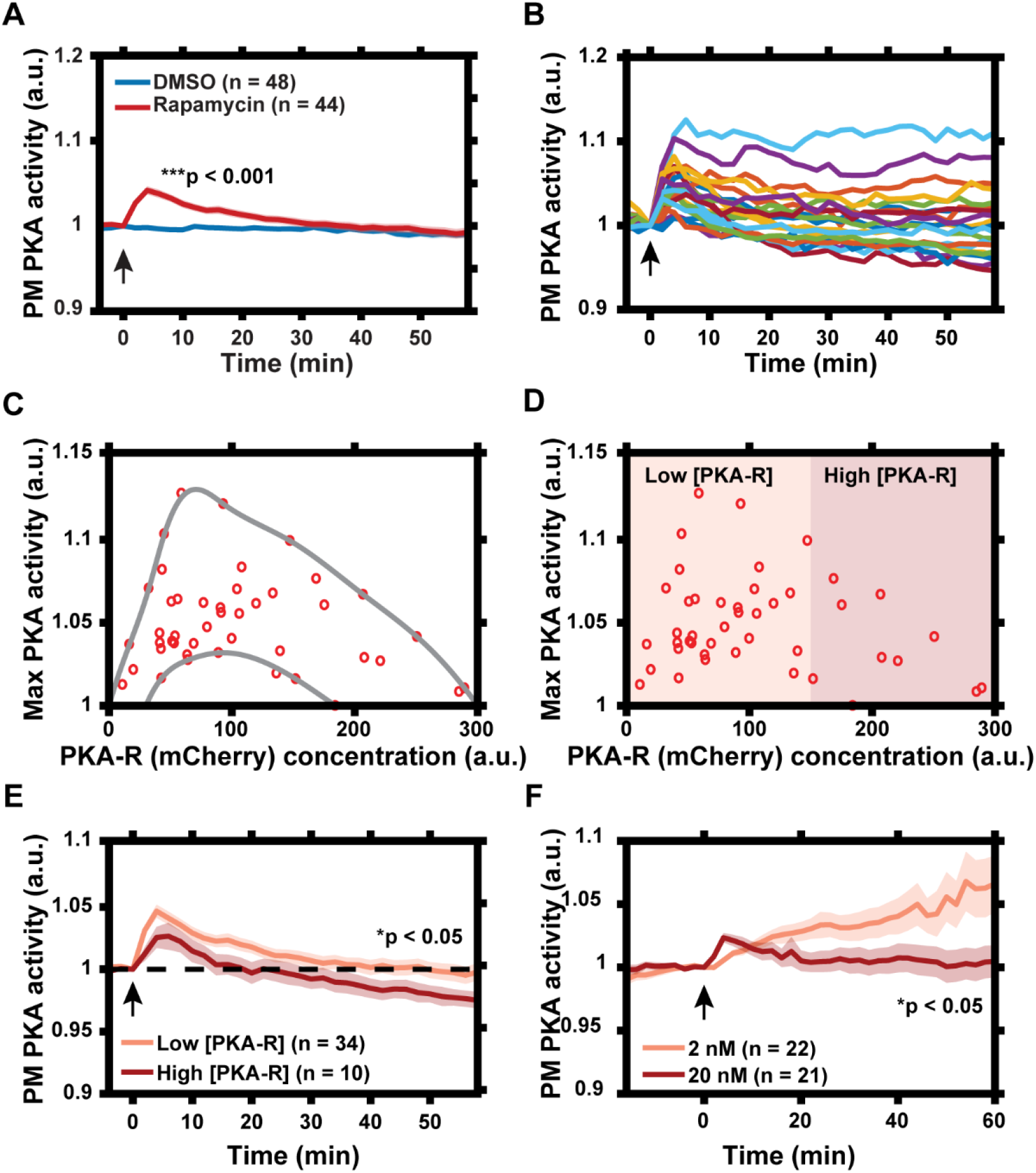
Characterization of PKA activity response to PKA-R translocation. (A) Effect of rapamycin-induced PKA-R translocation on PKA activity at the plasma membrane as detected by Lyn-AKAR4. 100nM rapamycin or 0.1% DMSO added at time = 0. p = 9.83 x 10^−12^ at t = 4 min post-rapamycin addition; two-tailed Student’s *t*-test. (B) Single cell PKA activity dynamics at the plasma membrane following addition of 100 nM rapamycin (a subset of data used for (A) is shown for clarity). (C) Relationship between PKA-R concentration, as estimated by mCherry fluorescence intensity, and maximal PM PKA activity increase following PKA-R translocation (n = 44 cells). Envelope overlaid for visualization. (D) Classification of cells into “low” and “high” expressors of PKA-R. (E) Average PM PKA activity over time for “low” versus “high” expressors of PKA-R (defined in panel D). p = 0.024 at t = 58 min post-rapamycin addition; two-tailed Student’s *t*-test. (F) PM PKA activity response to two different rapamycin doses. p = 0.023 at t = 60 min post-rapamycin addition; two-tailed Student’s *t*-test. Graphs in (A, E, F) display the mean of each data set with SEM indicated by shaded region. Number of cells in each data set is as indicated in the figure. Data is the result of 3 (A-E) and 2 (F) independent experiments, respectively. Arrows indicate the timing of drug addition.

To further investigate whether the local concentration of PKA-R played a role in determining the magnitude and duration of the response at the plasma membrane, we treated cells with two different lower concentrations of rapamycin – 2 nM and 20 nM. We again found that treatment of cells with 20 nM of rapamycin led to PKA activation that was, on average, transient, consistent with the gradual accumulation of PKA-R at the PM, first to optimal and then inhibitory levels. Importantly, we found that inducing a decreased level of PKA-R translocation with a lower dose of rapamycin (2 nM) resulted in a slower but much more sustained increase in PKA activity, reaching, on average, much higher levels than those seen for the higher dose (Figure 2F), suggesting that the lower PKA-R levels achieved at this rapamycin concentration were close to optimal. These results collectively suggested that PKA-R translocation to the PM can stimulate the PKA activity up to an optimal level of this subunit but can also inhibit PKA when the PKA-R levels exceed the optimal level in the PM compartment.

### Graded translocation of PKA-R induces a reversal of cell polarity

We next explored the functional effects of induced PM localization of PKA-R. PKA activity and PKA-R abundance have been shown to be elevated at the front of migrating cells *in vitro*, but the role that spatial localization of PKA plays in regulating cell migration is not well understood (Lim et al., 2008; Paulucci-Holthauzen et al., 2009). We therefore used our translocation system to probe the effect of intracellular PKA activity gradients, induced by gradients of PKA-R PM translocation, on cell polarization and migration. To accomplish this, we took advantage of the fact that intracellular PKA-R gradients can be induced by extracellular rapamycin gradients controlled within a microfluidic device (Lin et al., 2012; Lin & Levchenko, 2015; Lin, Yin, Wu, Inoue, & Levchenko, 2015). We used a variant of the microfluidic chip capable of generating chemical gradients by passive diffusion, which was previously developed in our lab (Lin et al., 2015). We increased the throughput to enable observation of up to 304 individual cells in parallel narrow microchannels (“cross-channels”) allowing for 1D cell migration (Figure 3A and Supplemental Methods). A gradient of rapamycin in the cross-channels was induced through diffusion between continuously replenished “source” and “sink” channels and visualized using a dye of similar molecular weight (Alexa Fluor® 594). The cells in the channels exposed to the gradient did not experience sheer stress because of a much higher hydraulic resistance within the channels relative to a much lower resistance in larger source and sink channels. HeLa PFY cells were seeded into the chip and allowed to migrate into cross-channels overnight. Due to directionality of cell seeding, most cells had an initial directional polarity of migration, resulting in the “upward” migration direction (from “sink” to “source,” as labeled in Figure 3A) before rapamycin was added.

**Figure 3.**
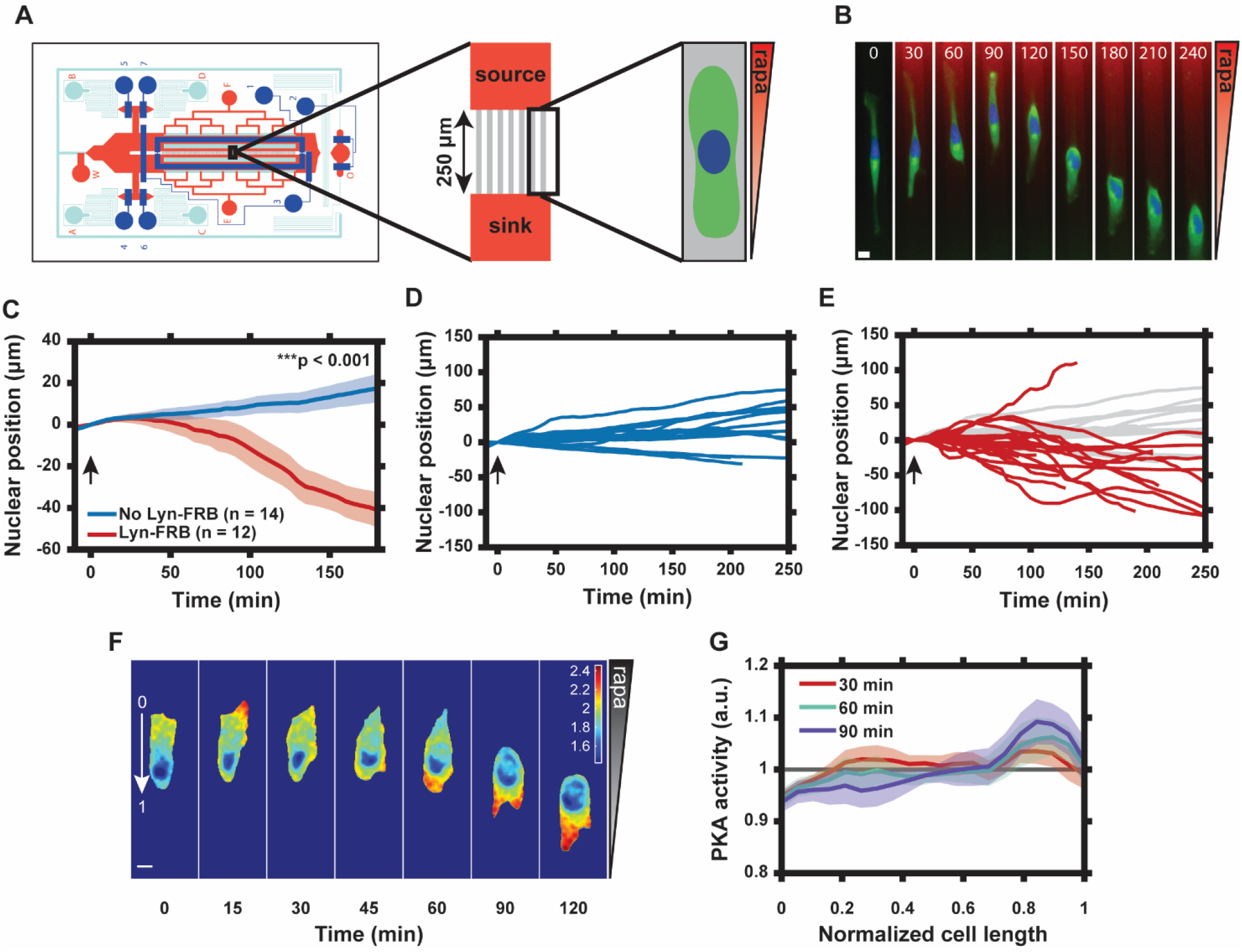
Graded translocation of PKA-R induces a reversal of cell polarity. (A) Schematic of the microfluidic device used to produce gradients of rapamycin across microchannels housing HeLa PFY or PFM cells. (B) Single cell response to 20 nM rapamycin gradient. Numbers show time in minutes. [Green = PKAR-FKBP-YFP, Blue = H2β-mCerulean (nuclear marker), Red = Alexa Fluor® 594 dye] (C) Average nuclear position (normalized to t = 0) for cells in 20 nM rapamycin gradient (0.08 nM/µm) versus no-translocation rapamycin control. SEM indicated by shaded regions. p = 1.34 x 10^−5^ at 180 min post-rapamycin addition; two-tailed Student’s *t*-test. Number of cells in each data set as indicated in the figure. (D) Single cell nuclear position data for cells in rapamycin gradient without Lyn-FRB. Data from 1 independent experiment. (E) Single cell nuclear position data for cells in 20 nM rapamycin gradient with Lyn-FRB. Data from 3 independent experiments. Data from (D) superimposed in light gray. (F) Tracking of intracellular PKA activity gradient with Lyn-AKAR4 following induction of rapamycin gradient at t = 0. Color indicates FRET ratio as in color bar. (G) Mean intracellular PM PKA activity tracking along the cell length from the high end of the rapamycin gradient (“0” in panel F) to the low end (“1” in panel F). Data represent the mean of n = 9 cells from 2 independent experiments with SEM indicated by shaded region. Cells were divided into 20 bins with the average FRET ratio value taken for each. Arrows in (C-E) indicate addition of rapamycin. Scale bars in (B) and (F), 10 µm.

Upon application of a rapamycin gradient (0 – 20 nM, 0.08 nM/μm), we found a pronounced reversal of the directionality of the cells’ migration in the direction opposite to the initial “upward” bias, with the preferred new direction thus being opposite to the direction of the rapamycin gradient (Figures 3C and 3E and Video 3). This was in stark contrast to cells without expression of Lyn-FRB or cells exposed to a DMSO gradient that did not experience PKA-R translocation and continued migrating in the “upward” direction (Figures 3C-3D, Figure 3 – figure supplement 1 and Video 4). This result was surprising and in apparent contradiction with the expectation that an increase in PKA-R at the cell front would enhance the preexisting cell polarization toward the rapamycin source. However, it was consistent with our prior observations suggesting that in many cells, the translocation of PKA-R induced by a sufficiently high rapamycin concentration could have an inhibitory rather than activating effect on PKA. This inhibitory effect was certainly true for a 20 nM spatially homogeneous dose of rapamycin, as revealed by the experiments and analysis described above (Figure 2F).

To determine whether PKA activity was indeed affected by a graded translocation of PKA-R, we repeated the experiment with HeLa PFM cells expressing Lyn-AKAR4. Prior to rapamycin exposure, most cells displayed an internal PKA activity gradient that corresponded with the cell polarization state (high PKA activity at the cell front and low at the rear) (Figure 3F). Upon exposure to the rapamycin gradient, we observed a reversal in the direction of the internal PKA activity gradient that was con-current with the flip in the direction of cell migration (Figures 3F and 3G). Interestingly, we also observed a transient increase in PKA activity at the cell front (Figure 3F, 15-minute timepoint) prior to this reversal. These results, in combination, further supported the explanation of the reversal of cell migration directionality due to a transiently positive but, in the longer run, negative effect of a high level of PKA-R PM translocation.

### Cell polarization state can be tuned by the slope of the rapamycin gradient

To further explore the mechanism of the reversal of cell migration directionality, we investigated how the slope and magnitude of the rapamycin gradient affected the response. First, we explored the effects of spatially uniform (20nM – 20 nM and 100nM – 100nM across the channel) rapamycin inputs. Both were expected to compete with internal polarity cues. We found that whereas the lower rapamycin concentration had no detectable effect (vs. e.g., the control Fig. 3D), the higher concentration gradually randomized the cell migration directionality, suggesting a stronger negating effect on the intrinsic polarity regulation (Fig. 4A). We then exposed the cells to a relatively shallow rapamycin gradient (10 nM – 20 nM across the channel, or 0.04 nM/μm), contrasting the results with those produced by steeper spatially graded stimulation (0 – 20 nM, or 0.08 nM/μm, Figure 3E). We again found reversal of the average migration directionality, which was increasingly more pronounced with increasing gradient slope (Figures 4D and 4F). This result suggested that both gradient values were sufficient to compete with the intrinsic polarity regulation mechanisms, by suppressing PKA activity in the front of the cell greater than in the rear. By this logic, we expected that presenting cells with a reverse gradient (20 – 0 nM, “downward”) would result in internal gradients of PKA activity that would effectively point “upward” for a large subset of cells, which would be consistent with the directionality of their inherent polarity, an effect opposite to the rapamycin gradients pointing “upward”. Thus, cells were expected to continue moving “upward”, perhaps at an even greater rate vs. the control. We indeed observed that most cells under this condition moved similarly to the control case, while a subset of cells reversed their directionality from “downward” to “upward”, and another subset displayed an increased speed of “upward” migration vs. the maximal cell migration levels observed in the control (Figures 4G and 4H).

**Figure 4.**
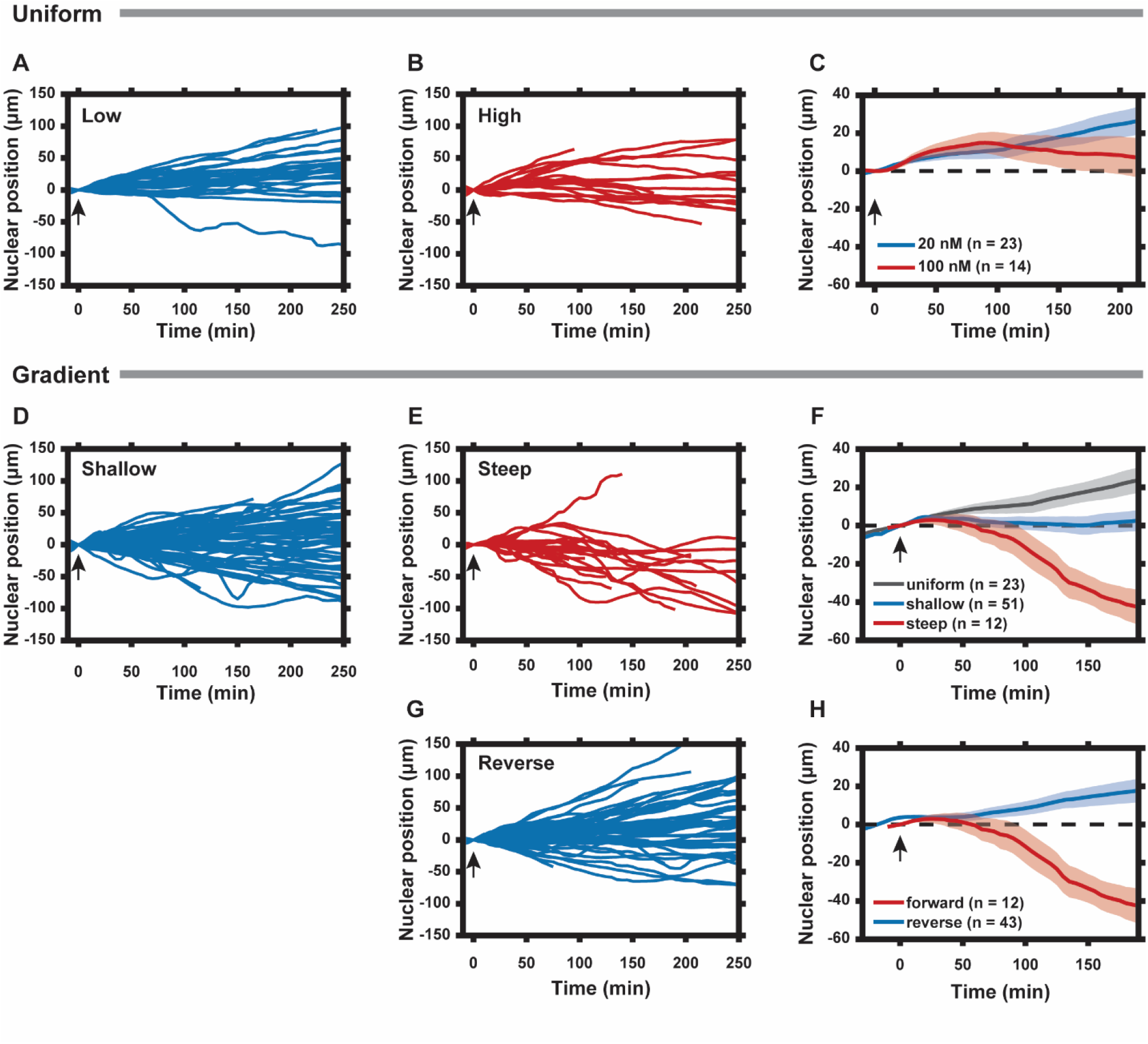
Cell polarization state can be tuned by the slope of the PKA-R gradient. (A) Nuclear positions of single cells responding to a uniform 20 nM rapamycin stimulus. (B) Nuclear positions of single cells responding to a uniform 100 nM rapamycin stimulus. (C) Average nuclear position of cells responding to 20 nM versus 100 nM uniform rapamycin stimulus. SEM indicated by shaded regions. (D) Nuclear positions of single cells responding to a shallow rapamycin gradient (10 nM - 20 nM, or 0.04 nM/µm). (E) Nuclear positions of single cells responding to a steep rapamycin gradient (0 nM - 20 nM, or 0.08 nM/µm). Data taken from Fig. 3E. (F) Average nuclear position of cells responding to a uniform 20 nM stimulus from (A), shallow gradient from (D), or steep gradient from (E). SEM indicated by shaded regions. (G) Nuclear positions of single cells responding to a steep rapamycin gradient in the reverse direction (20 nM - 0 nM, or 0.08 nM/µm) as what is shown in (3A). (H) Average nuclear position of cells responding to steep rapamycin gradient in the forward and reverse directions. SEM indicated by shaded regions. Arrows indicate addition of rapamycin. Number of cells in (C), (F), and (H) are as indicated in the figure. Data was collected from 2 (A, D, G) and 1 (B) independent experiments.

## Discussion

Proper kinase localization is important for cell function, ensuring that the kinase activity is limited to a specific set of substrates in response to an extracellular cue. Here, we describe development and implementation of a novel approach to dynamically relocalize a regulatory subunit of a kinase, PKA, to the plasma membrane. In contrast to a previous relocalization technique utilizing a photoactivatable PKA-C, our approach maintains the dependence of resulting PKA activity on cAMP and does not require overexpression of the catalytic subunit (O’Banion et al., 2018). Furthermore, our synthetic system recapitulates the natural mechanism of subcellular PKA targeting of the regulatory PKA subunits by AKAPs.

In regulation of PKA activity, the regulatory PKA subunit can play a dual role of an inhibitor of PKA-C and a mediator of PKA localization to specific subcellular compartments, potentially enriched in the kinase activator (cAMP) and kinase substrates. Thus, in spite of its canonically inhibitory function, PKA-R can potentially have a more positive, up-regulating role on the activity of the enzyme. Our approach has allowed us to clarify this paradoxical function of PKA-R. We demonstrate in particular that relocalization of PKA-R to the plasma membrane was sufficient to induce an increase in PKA activity in this intracellular compartment, particularly for lower levels of the translocated PKA-R, even in the absence of changes in upstream signaling through G protein-coupled receptors. The effect of the PKA-R translocation reached the maximal level at the optimum level of this subunit, decreasing at PKA-R higher levels and ultimately driving the PKA activity to below basal levels. This type of nonlinear behavior has been experimentally and computationally demonstrated for linker proteins, such as scaffold proteins (Chapman & Asthagiri, 2009; Good et al., 2011; Levchenko et al., 2000). At low levels, a scaffold can enhance signaling by bringing components of a signaling pathway into close proximity, enabling their interaction, and thus enhancing the pathway activity. However, when the scaffold concentration is too high, it sequesters pathway components away from each other, decreasing the pathway activity. Our results suggest that within the cell, PKA-R plays a scaffold-like role for PKA-C and cAMP since the kinase activity requires linkage of two PKA-C subunits and four cAMP molecules into one molecular complex by these subunits. When the local PKA-R concentration is too high, it can lead to incomplete binding (titration away) of either cAMP or PKA-C, preventing activation. Therefore, as PKA-R is recruited to the membrane, PKA activity will increase until there is a local shortage of one or both of these components. This suggests a more complex view of PKA-R’s role in this pathway, suggesting that it can have either a pathway inhibitory or enhancing role, depending on the expression of PKA-C or cAMP abundance both globally across the cell, and in specific subcellular compartments. In particular, our results suggest that the abundance of PKA-R at the plasma membrane in the HeLa cells investigated here is suboptimal, but one can expect that it can be optimal or ‘super-optimal’ in other cellular compartments.

Our tool also allowed us to study the functional effect of controlling the basal PKA activity on a complex phenotypic response: the polarity of cell migration. Although PKA-R has been shown to be enriched at the leading edge (Howe et al., 2005) of moving cells, the functional importance of either this localization pattern or the overall PKA activity gradient across a polarized cell has not been fully understood. Indeed, PKA activity gradients can be triggered by a number of membrane localized receptors or other signaling proteins, which are frequently pleiotropic and can activate other signaling pathways. Within this in vivo complexity, our molecular tool can isolate the specific role of PKA activity in defining cell polarity. It can be particularly revealing if one can reverse the cell polarity by directly imposing an intracellular PKA activity gradient. We indeed found that inducing gradients of PKA-R PM translocation at levels inhibitory to PKA activity led to a reversal of the internal PKA activity gradient and, as a result, the cell polarity and direction of cell migration. This work therefore demonstrates for the first time that intracellular PKA activity gradients can provide a polarization cue that is powerful enough to overcome other existing polarization cues inside the cell.

It is likely that PKA activity gradients exert an effect on cell polarization via modulation of the activity of the two mutually inhibitory Rho-family small GTPases RhoA and Rac1. PKA phosphorylation of RhoA on Ser188 has been shown to promote its interaction with RhoGDI, leading to removal of RhoA from the PM (Lang et al., 1996). PKA is also implicated in Rac1 activation, although the direct target of this positive regulation is still unclear (O’Connor & Mercurio, 2001). Since PKA has a positive effect on Rac1 activity and a negative effect on membrane localized RhoA activity, it would be reasonable to theorize that PKA acts to shift the balance of Rac1 to RhoA signaling in favor of Rac1. In this way, a gradient of intracellular PKA activity could help to enhance the internal polarization of these two polarity effectors. When the PKA activity gradient flips, as in Figure 3G, the effect would be to increase Rac1 and decrease RhoA activity at the cell rear, leading to a reversal of polarity if this signal is sufficiently strong.

In a broader sense, this work demonstrates the critical role that kinase localization plays in controlling the cell response to an incoming stimulus. The subcellular kinase concentration can be tuned by the expression levels of various scaffold proteins, and differential expression between cells may result in variability of signaling responses within a population. Localization of scaffold proteins may vary dynamically via modifications to subcellular localization sequences as well, resulting in similar effects to those described here. In the context of PKA, AKAPs are receiving attention as potential therapeutic targets, and at least one isoform of PKA-R has been implicated as a driver of oncogenesis (Codina et al., 2019; Esseltine & Scott, 2013; Veugelers et al., 2004). The kinase relocalization strategy demonstrated herein can be applied to study the effects of localized PKA activity in other subcellular compartments and can be adapted to the study of other kinases, enabling a better understanding of the role of compartmentalized signaling.

## Materials and Methods

### Cell culture and reagents

HeLa cells were cultured in high glucose Dulbecco’s modified Eagle medium (DMEM, Corning, Corning, NY) supplemented with 10% fetal bovine serum (ThermoFisher Scientific, Waltham, MA) and 1% penicillin/streptomycin (ThermoFisher Scientific). For stable lines, media was supplemented with 1µg/ml puromycin (MilliporeSigma, Burlington, MA) and blasticidin (ThermoFisher Scientific) to continuously select for cells expressing PKAR-FKBP-FP and Lyn_11_-FRB (also referred throughout to as Lyn-FRB). Cells were cultured in a 37°C humidified incubator with 5% CO_2_. Rapamycin was obtained from LC Laboratories. Forskolin and IBMX were obtained from MilliporeSigma. The Lyn-AKAR4 FRET probe was a gift from Dr. Jin Zhang (University of California San Diego). Transient transfections were completed using Fugene (Promega, Madison, WI) according to manufacturer’s protocol. Transfection of lentiviral vectors was completed using TurboFect (ThermoFisher Scientific).

### Plasmid design

RNA was isolated from HeLa cell lysate and PCR amplified for PKA-RIIβ and PKA-Cβ respectively using custom primers. The gel purified PCR product was ligated into a transient expression plasmid containing a YFP or mCherry fluorescent protein before being transformed into DH5α competent cells. Amplified plasmid was recovered using a standard mini prep kit (Qiagen, Hilden, Germany) and sent for sequencing.

### Cell line generation

PKAR-FKBP-FP and Lyn-FRB were transferred to lentiviral expression vectors by Gateway cloning. Both color variants of PKAR-FKBP-FP were cloned into pLenti CMV puro destination vectors (Addgene plasmid #17452). Lyn-FRB was cloned into the pLenti CMV blast destination vector (Addgene plasmid #17451). To generate lentivirus, HEK293FT cells were transfected with one of the three destination vectors plus a lentiviral packaging vector (psPAX2, Addgene plasmid #12260) and a VSV-G envelope expressing vector (pMD2.G, Addgene plasmid #12259). Cell media was collected over a three-day period beginning two days post-transfection and spun down in order to collect supernatant. Then, lentivirus was recovered from supernatant using an Amicon Ultra Centrifugal 50kDa filter (MilliporeSigma) and transduced into HeLa cells along with 10 μg/ml polybrene (Santa Cruz Biotechnology, Dallas, TX). Cells were transduced sequentially with PKAR-FKBP-FP followed by Lyn-FRB. Each time, a selection procedure was completed, using 10 μg/ml puromycin or blasticidin respectively, followed by generation of clonal lines by limiting dilution. Following selection, stable cells were cultured in DMEM media containing 1µg/ml puromycin and blasticidin to maintain expression of the PKA-R translocation system.

### Microfluidic device fabrication

Multilayer microfluidic devices were fabricated from polydimethylsiloxane (PDMS) via replica molding from custom silicon masters, as described in Supplemental Methods. Devices were cleaned with 2% Alconox and 70% ethanol and then thermally bonded to #1.5 glass coverslips (Corning) by a 24-hour bake at 85°C. See Supplemental Methods for further details.

### Microfluidic device setup

Microfluidic channels were coated with 10 μg/ml fibronectin by incubation for 1 hour at room temperature. Cells were suspended in imaging medium (DMEM without phenol red, 10% FBS, 1% pen-strep) and seeded into the device as described in Supplemental Methods. Following overnight incubation at 37°C, a microfluidic valve was depressed to fluidically separate the cells from all future inputs and imaging medium containing rapamycin (or an equivalent volume of DMSO) was added to one or both sides of the device. A gradient from one side of the microchannels to the other was established by creating a height-driven pressure differential between two input and one output ports. Following the start of the imaging, the microfluidic valve was gradually released to expose cells to the gradient.

### Imaging

Widefield imaging was performed on a Zeiss Axiovert 200 M epifluorescence microscope with motorized stage (Prior, Rockland, MA) and live cell incubation chamber with humidity and temperature control (PeCon, Erbach, Germany) set to 37°C and 5% CO_2_. Cells were imaged using a 40X, 1.3 numerical aperture oil immersion objective (Zeiss, Oberkochen, Germany) and Cascade II:1024 EMCCD camera (Teledyne Photometrics, Tucson, AZ). Microscope automation was controlled with Slidebook 6.0 software (Intelligent Imaging Innovations, Denver, CO). PKA-R was imaged using YFP excitation and emission filters for HeLa-PFY cells or mCherry excitation and emission filters for HeLa-PFM cells. For nuclear tracking experiments, cells were transfected with H2β-mCerulean, which was detected using CFP excitation and emission filters. All FRET images were obtained using a CFP excitation filter as well as CFP and YFP emission filters. Semrock filters and corresponding dichroics were used (IDEX Health & Science, LLC, Rochester, NY). For the migration experiments, a spectral 2D template autofocus algorithm was employed using the phase channel to correct for focus drift between timepoints.

Confocal imaging was performed on a Nikon TiE inverted microscope (Nikon, Tokyo, Japan) equipped with a Yokogawa CSU-W1 spinning disk with 50µm disk pattern (Yokogawa Electric, Tokyo, Japan) and CFI Plan Fluor 40X, 1.3 numerical aperture oil immersion objective (Nikon). Images were captured using an Andor iXon Ultra888 EMCCD camera (Oxford Instruments, Abingdon, UK). The microscope was equipped with a stage top incubator (Okolab, Pozzuoli, NA, Italy) maintaining humidity and 5% CO2.

### Live cell imaging

For dish experiments, 35 mm glass bottom dishes #1.5 (Matsunami, Osaka, Japan) were coated with 5-10µg/ml fibronectin (MilliporeSigma) for 1 hour at room temperature, and cells were incubated on the coated surfaces overnight prior to imaging. For FRET analysis, cells were imaged in Hank’s balanced salt solution (ThermoFisher Scientific) to reduce background. Other imaging experiments were completed in cell culture medium without phenol red. For analysis of translocation, images were taken every minute in the YFP and/or mCherry channels. For FRET, images were taken every 2 minutes in the CFP and FRET channels.

### FRET image analysis

All image analysis was performed using custom MATLAB codes. For FRET images, image registration was performed prior to analysis. Images were segmented by intensity thresholding in the YFP channel. The FRET ratio was calculated as (FRET-DF)/(CFP-DF), where FRET is the intensity of emission collected in the YFP channel following CFP excitation. Cells were excluded from the analysis if the expression of the FRET probe was below a threshold value (as determined by fluorescence intensity) that was consistent across experimental replicates.

### Cytoplasmic intensity tracking

To verify translocation of PKA-R and PKA-C to the plasma membrane, the cytoplasmic intensity of fluorescently tagged PKA-R or PKA-C was tracked over time. This was done by: (1) segmenting the cells in the mCherry channel (for PKA-R tracking) or YFP channel (for PKA-C tracking) respectively; (2) eroding away the outermost 20 pixels (7.78 μm) of the segmented cell, leaving only the fluorescence intensity values for the non-plasma membrane compartments; and (3) taking an average of the non-zero pixels at each timepoint in the YFP channel (PKA-R) or mCherry channel (PKA-C) respectively. This average was tracked as a function of time before and after rapamycin stimulation. Cells were excluded from the analysis if the intensity in the fluorescence channel used for segmentation was too low to properly distinguish cell from background.

### Cell migration analysis by nuclear tracking

Prior to seeding into microfluidic devices, cells were transfected with H2β-mCerulean for nuclear visualization and tracking. Following experimentation, images were registered in the phase channel to prevent any miscalculations of nuclear position due to stage drift. Then, the nucleus was identified by intensity thresholding in the CFP channel, the centroid of each nuclear ROI was calculated, and its location was tracked over time. Since cells were confined to motion in 1 dimension and devices were aligned such that microchannels were parallel to the stage mount, only the y position of the centroid was required to capture the change in nuclear position with respect to time zero. This positive or negative change in nuclear position was graphically presented for both single cells and as an average of cells in a population.

### Statistical Analysis

Experimental results were presented either as single-cell traces or as means with shaded regions indicating SEM. Numbers of biological and technical replicates are as indicated in each figure. A biological replicate is defined as a single cell in an imaging experiment. A technical replicate is defined as a single imaging experiment. Each technical replicate is performed in a different cell imaging dish or microfluidic chip. Statistical analysis was completed in Microsoft Excel. Comparisons between groups were conducted using a two-tailed Student’s t-test. Comparisons were concluded to be significant for P values less than 0.05.

## Acknowledgements

We thank Dr. Jin Zhang (UCSD) for the gift of FRET probe (Lyn-AKAR4). We are grateful to Yegor Isakov for assistance in creating PKAR-FKBP-FP and PKAC-mCh transient expression plasmids. We are grateful to Drs. Eric Chu and Hao Chang for the assistance in microfluidic fabrication and molecular biology techniques.

## Supplemental Methods

### Design of the Microfluidic Device for the Analysis of the effect of PKA-R gradients on Cell Migration

To recapitulate the PKA gradients that have been seen in vitro, we needed a means of generating precise gradients of rapamycin. To do this, we utilized a diffusion-based microfluidic gradient generator (Supplementary Figure 1) that is based on a previous design from the Levchenko lab (Lin & Levchenko, 2015; Lin et al., 2015). This device can generate soluble gradients that are stable over the time scale of our experiments (> 6 hours). Briefly, a gradient is established within 6-μm tall parallel microchannels (gray) by passive diffusion from 130-μm tall perpendicular source and sink channels (light blue). The medium (and drug) is continually replenished by flow, which is driven by a hydrostatic pressure difference from two inlets (A and B or C and D) to a single outlet (O). Temporal control of experimental inputs is provided by a series of valves (dark blue) that, when depressed, can cut off flow to the microchannels. This two-layer design with valves was pioneered by the Quake group (Unger, Chou, Thorsen, Scherer, & Quake, 2000). Shear stress on cells seeded into the microchannels is minimized due to the large height difference between the source and sink and the microchannels, resulting in a much higher resistance in the microchannels and ensuring that the primary means of solute transport to the cells is diffusion. The microchannels are 16-μm wide, forcing the cells to assume a uniaxial phenotype. This reduces any analysis of cell migration to one dimension while providing a realistically confined environment. We increased the number of microchannels by over 20% from the original design (from 250 to 304) to increase the number of single cells that can be observed in parallel.

**Supplementary Figure 1.**
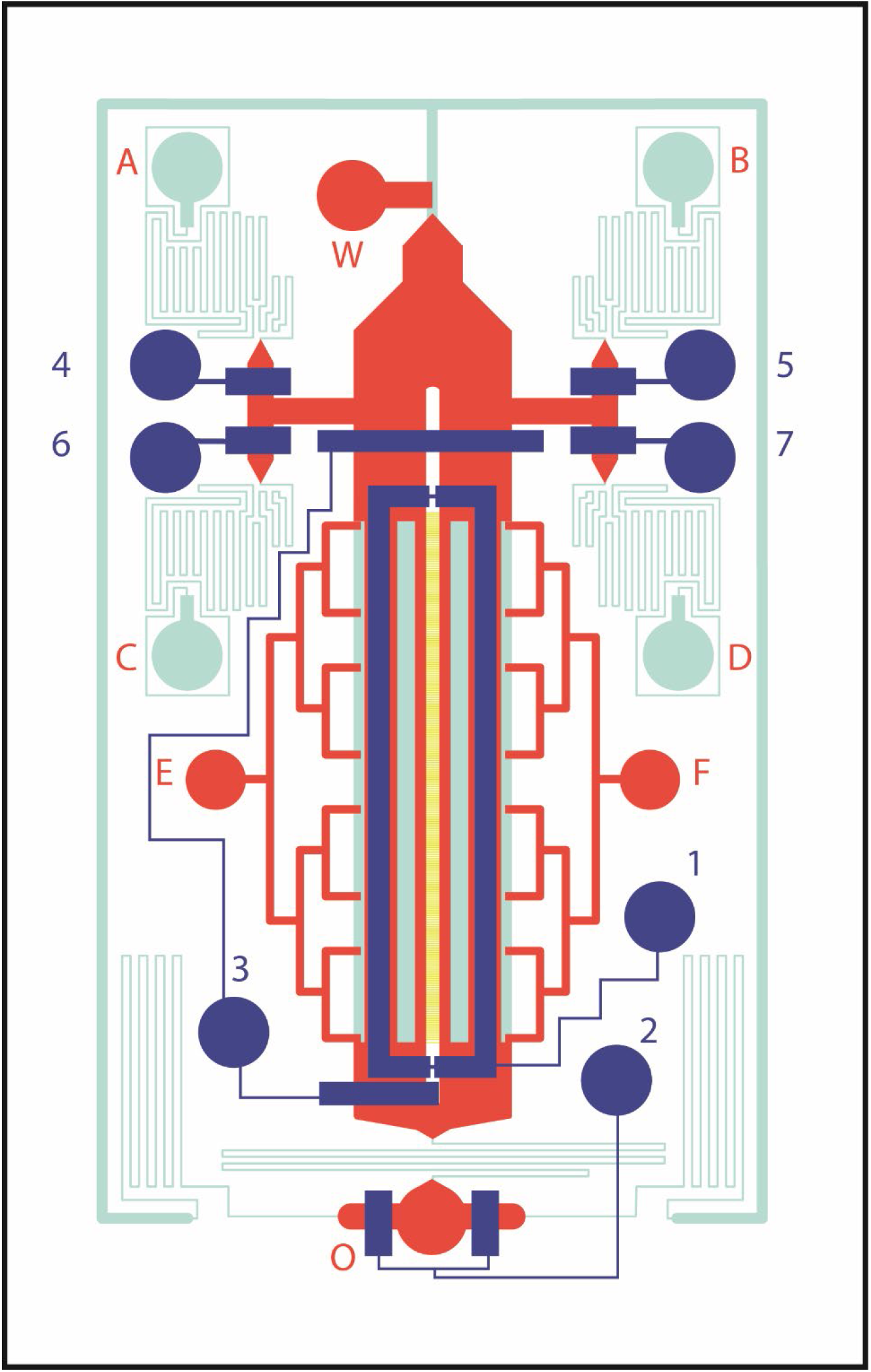
Microfluidic gradient chip used in the analysis. Cells are seeded into 16 µm wide x 6 µm tall microchannels (yellow) with gradient supplied by passive diffusion from adjacent 130 µm tall source and sink channels (light blue). Valves (dark blue) provide pressure to depress underlying 30 µm tall rounded fluidic channels (red). During experiment, media (with or without drug) flows from inlet ports A/B or C/D to outlet port O.

### Microfluidic device fabrication

Microfluidic devices were fabricated from polydimethylsiloxane (PDMS) via replica molding from silicon masters. The silicon masters were fabricated in a clean room using a process known as photolithography in which photo-sensitive resists are spin-coated onto silicon wafers and regions corresponding to desired features are selectively cured with UV light (Sia & Whitesides, 2003). Photomasks for each master were designed using Adobe Freehand software and printed as transparencies.

Since our device consists of 2 layers of PDMS (a flow layer and an overlying valve, or control, layer), two separate masters were prepared. For the control layer, SU-8 2025 (Microchem) was deposited onto a 3” silicon wafer and spin-coated according to manufacturer’s protocol to achieve a desired feature height of 40 μm. Features were selectively exposed to UV light in a mask aligner with a constant energy of 1000 mJ/cm^2^. Following exposure, the SU-8 was baked at 95°C for 1 hour followed by removal of uncured resist with SU-8 developer (Microchem) and a second bake at 200°C for 1 hour to reinforce the features.

For the second master, a similar protocol was followed. The flow layer includes features of 3 different heights: 6 μm, 30 μm and 130 μm. Thus, the preparation of the master was conducted in three stages. SU-8 2005 (Microchem) was spin-coated onto a 3” silicon wafer to form the 6 μm level. Then, SPR-220-7 resist (Megposit) was spun onto the wafer at 1000 rpm for 30 seconds to form the 30-μm level Finally, SU-8 2025 (Microchem) was spun onto the wafer at 1500 rpm according to manufacturer’s protocol to form the 130-μm level. In between each spin-coating step, standard photolithography procedures were followed for UV exposure and baking of the channel regions as outlined above. For the SPR layer, a special hard bake protocol was followed to create rounded channels that could be depressed by pressurizing the overlying valves. Specifically, the wafer was baked for 5 hours while ramping the temperature at a rate of 180°C per hour to a final temperature of 200°C.

PDMS devices were fabricated by replica molding from the silicon masters. First, masters were functionalized with 1H,1H,2H,2H-Perfluorooctyltrichlorosilane overnight to facilitate removal of PDMS from the mold. Then, PDMS (RTV615, Momentive) was prepared in a 20:1 ratio for the flow layer and 5:1 ratio for the control layer. Following initial degassing, PDMS was poured over the flow layer wafer and placed in a vacuum chamber for an additional 15 minutes to ensure that no air bubbles remained in the channel regions. The wafer was then spin coated for 1 minute at 390 rpm to achieve a uniform, thin layer of PDMS and then baked at 70°C for 15 minutes until partially cured. For the control layer, PDMS was poured over the silanized master and baked for 40 minutes at 70°C. Then, the valves were cut out, inlets were added using a 20-gauge luer stub, and valves were aligned on top of the flow layer. The two layers were then cross-linked to one another by baking at 70°C for at least 2 hours. Complete devices were cut from the silicon master and inlets added as before. Chips were cleaned with a combination of tape, 2% Alconox solution, and 70% ethanol prior to thermal bonding to #1.5 glass coverslips (Corning) at 85°C for 24 hours.

### Microfluidic device setup

Control valves 1-3 (Figure 3) were filled with DI water using 10 PSI pressure driven by a set of solenoid valves (the Lee Company). Then, the pressure to valve 1 was released, and valves 2 and 3 were pressurized to ∼20 PSI to block flow from passing between the microchannel region and the rest of the fluidic network. A syringe containing 10 μg/ml fibronectin was inserted via syringe tubing into port E and elevated to force flow through the microchannels to ports F and O, which were then blocked.

Following 1 hour of fibronectin coating, the fibronectin was rinsed out with imaging medium (DMEM without phenol red, 10% FBS, 1% pen-strep) through a syringe at port F. The rest of the fluidic network was filled through temporary release of valve 2 and insertion of syringes with imaging medium at either ports A and B or C and D. All other ports were blocked.

Next, a cell suspension was added as a droplet at port E, and cells were flowed into the microchannel region via negative pressure provided by lowering the syringe at port F. Once sufficient cells had been added and were lining the microchannels, valve 1 was pressurized to avoid disturbing the cells with future manipulations. The syringe at port F was then removed and a fresh media syringe was added at port O. After blocking all ports, the valves were slowly released, and the device was transported to a 37°C incubator with 5% CO_2_. Care was taken to ensure that all 3 syringes were kept at equal heights to prevent flow through the device. After 3 hours, the syringe in port O was lowered by 1 inch to maintain low velocity flow of fresh media during overnight incubation.

Following overnight incubation, valve 3 was depressed to isolate cells from future manipulations. Then, one or both syringes at the top of the device were replaced with media containing rapamycin and Alexa Fluor® 594 dye to track the development of the gradient (ThermoFisher). The device was then transferred to the microscope, and a 2.5-inch pressure difference was established between the inlet and outlet ports to drive fluid flow and maintain the gradient. Following the start of imaging, valve 3 was slowly released to expose cells to the rapamycin gradient.

## Figure Supplements

**Figure 1 – figure supplement 1.**
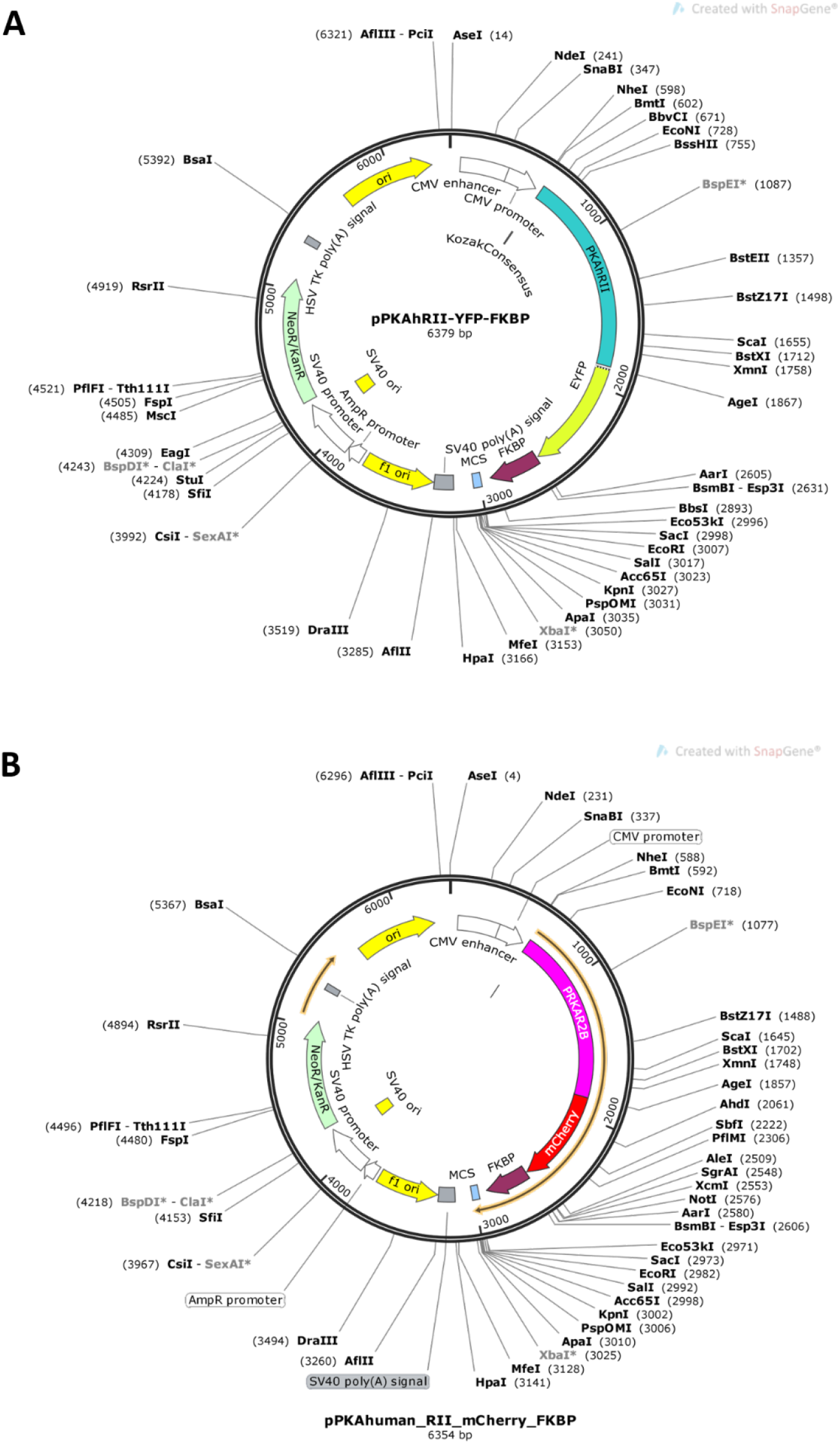
Plasmid maps for PKAR-FKBP-FP constructs. Plasmid maps for (A) PKAR-FKBP-YFP and (B) PKAR-FKBP-mCherry.

**Figure 2 – figure supplement 1.**
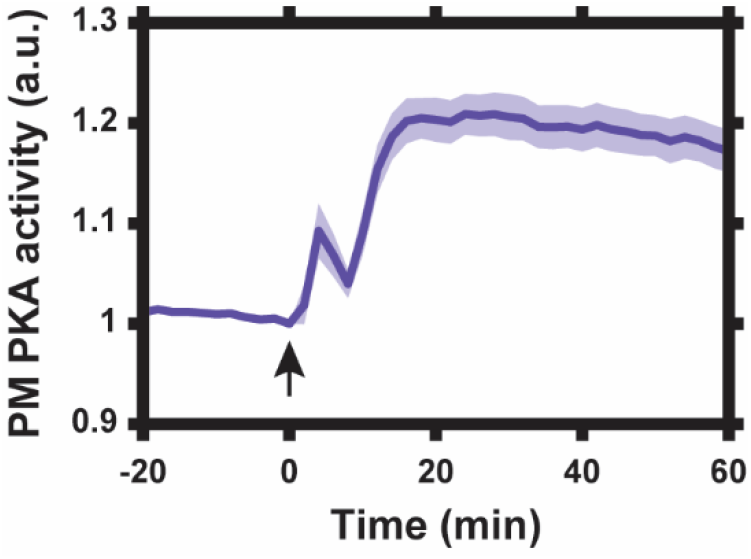
Lyn-AKAR4 dynamic range. Lyn-AKAR4 FRET response in HeLa cells following co-treatment with 50 μM forskolin and 100 μM IBMX to maximally increase intracellular cAMP levels. Data represent the mean of n = 12 cells from 1 independent experiment with SEM indicated by shaded region. Arrow indicates drug addition.

**Figure 3 – figure supplement 1.**
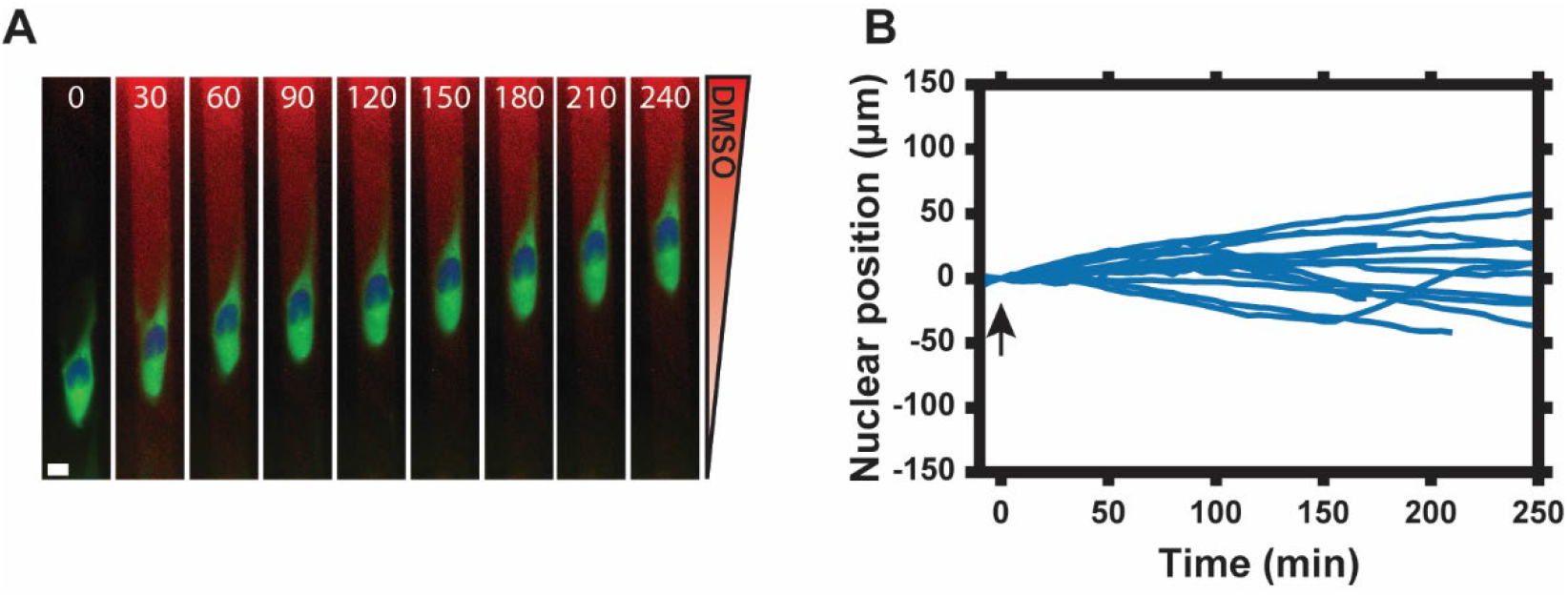
1D cell migration in DMSO gradient. Single cell response to DMSO gradient (0 – 0.1% from “sink” to “source” as labeled in Figure 3.3A). Numbers show time in minutes. Scale bar, 10 µm. [Green = PKAR-FKBP-YFP, Blue = H2β-mCerulean (nuclear marker), Red = Alexa Fluor® 594 dye] (B) Nuclear position normalized to t = 0 min post-DMSO addition. A positive nuclear position indicates net movement of the nuclear centroid toward the DMSO source as described in Materials and Methods. Arrow indicates DMSO addition. Data represent nuclear position for n = 18 cells from 2 independent experiments.

## Video Descriptions

***Video 1. Rapamycin-induced translocation of PKA-R***. Translocation of PKA-R (green) to the plasma membrane of HeLa cells following treatment with 100 nM rapamycin at time zero as indicated in the video. Cells were co-transfected with mCherry protein (red) as a counterstain for visualization.

***Video 2. Co-recruitment of PKA-C and PKA-R to the plasma membrane***. Colocalization of PKA-C (red) and PKA-R (green) before and after treatment with 100 nM rapamycin at time zero as indicated in the video.

***Video 3. Single cell response to graded PKA-R recruitment***. Single cell response to 20 nM rapamycin gradient applied at time zero as indicated in the video. [Green = PKAR-FKBP-YFP, Blue = H2β-mCerulean (nuclear marker), Red = Alexa Fluor® 594 dye]

***Video 4. Single cell response to DMSO gradient***. Single cell response to DMSO gradient applied at time zero as indicated in the video. [Green = PKAR-FKBP-YFP, Blue = H2β-mCerulean (nuclear marker), Red = Alexa Fluor® 594 dye]

## Source Data

**Figure 1 – source data 1**

**Characterization of PKA-R translocation system**

(a) Sheet 1, Figure 1D Time Course. Cytoplasmic intensity change in the YFP channel following addition of 0.1% DMSO or 100 nM rapamycin. Mean, SEM, and number of cells given for each timepoint and condition. (b) Sheet 2, Figure 1E Time Course. Cytoplasmic intensity change in the mCherry channel following addition of 100 nM rapamycin. Mean, SEM, and number of cells given for each timepoint and condition.

**Figure 2 – source data 1**

**Lyn-AKAR4 data following PKA-R translocation**

(a) Sheet 1, Figure 2A Time Course. Lyn-AKAR4 response following addition of 0.1% DMSO or 100 nM rapamycin. Mean, SEM, and number of cells given for each timepoint and condition. (b) Sheet 2, Single cell data. Single cell time courses from Figures 2A-2E. Cells are labeled as having low or high PKA-R expression. (c) Sheet 3, Figure 2E Time Course. Lyn-AKAR4 response following addition of 100 nM rapamycin for cells with low versus high PKA-R expression. Mean, SEM, and number of cells given for each timepoint and condition. (d) Sheet 4, Figure 2F Time Course. Lyn-AKAR4 response following addition of 2 nM or 20 nM rapamycin. Mean, SEM, and number of cells given for each timepoint and condition.

**Figure 3 – source data 1**

**Nuclear position and Lyn-AKAR4 data in microfluidic device**

(a) Sheet 1, Figure 3C and Figure 3 – figure supplement 1 Time Course. Normalized nuclear position data for cells in 20 nM rapamycin gradient (0.08 nM/µm) with or without the membrane subunit of the PKA-R translocation system (Lyn-FRB). Results also shown for cells in a volume equivalent DMSO gradient, as displayed in figure supplement 1. Mean, SEM, and number of cells given for each timepoint and condition. (b) Sheet 2, Figure 3G Intracellular PKA Activity Distribution. Mean intracellular PM PKA activity along the cell length from the high end of the rapamycin gradient (“0”) to the low end (“1”) for 3 time points. Mean, SEM, and number of cells given for all 20 bins and each timepoint.

**Figure 4 – source data 1**

**Nuclear position data for cells exposed to different rapamycin stimuli in microfluidic device**.

Figure 4 Time Courses. Normalized nuclear position data for cells exposed to rapamycin distributions of varied magnitudes, slopes, and orientations. Mean, SEM, and number of cells given for each timepoint and condition.

